# Simultaneous Optimization of Codon Adaptation and Thermodynamic Stability in mRNA Therapeutics via Deep Reinforcement Learning

**DOI:** 10.64898/2026.11.26.769446

**Authors:** Baisunkar Mirasbek, Martin Eberhardt, Carola Berking, Jan Dörrie, Julio Vera

**Affiliations:** Department of Dermatology, FAU Erlangen-Nürnberg, Erlangen, Germany; Hautklinik, Uniklinikum Erlangen, Erlangen, Germany; BZKF, Erlangen, Germany; CCC Erlangen-EMN, Erlangen, Germany; Deutsches Zentrum Immuntherapie (DZI), Erlangen, Germany

**Keywords:** mRNA vaccine design, codon optimization, deep reinforcement learning, RNA’s minimum free energy, codon adaptation index, multi-epitope vaccine, multi-objective optimization

## Abstract

**Motivation:** The efficacy of mRNA vaccines depends on translational efficiency and molecule stability, commonly captured by the codon adaptation index (CAI) and minimum free energy (MFE), respectively. These objectives are in tension, and codon degeneracy makes their simultaneous optimization in large RNA constructs encoding multiples peptides a combinatorial search problem intractable through brute-force computation.

**Methods:** We present ReinforSeqPipe, a Deep Q-Network reinforcement learning pipeline coupling a Long Short-Term Memory (LSTM) sequence encoder with the bi-parametric MFECAIλ objective as its reward function. Adapted from LinearDesign, MFECAIλ is a weighted objective function of MFE- and CAI-centred subfunctions. Operating without training data, the agent performs synonymous mutagenesis guided solely by the biophysical reward, with parallel stochastic runs enabling best-of-N selection. The pipeline was applied to all 120 permutations of a five-epitope HLA-A*02:01-targeting construct at λ ∈ {1, 4, 10}, with λ being a parameter mediating the MFE–CAI trade-off (low λ prioritizing MFE and high λ prioritizing CAI). The best RNA designs were validated experimentally in Jurkat cells.

**Results and Conclusions:** Optimized sequences achieved significantly higher MFECAIλ than codon-optimized controls (145.6 ± 5.8 vs. 109 ± 1.8, p < 0.001), driven predominantly by improved MFE at modest CAI cost. Within ten stochastic repeats, ReinforSeqPipe matched or surpassed LinearDesign at every λ setting, leading on the majority to all 120 permutations depending on λ. Experimentally, the optimized mRNA showed a significantly longer half-life than synonymous and non-synonymous controls in Jurkat cells (∼20-minute increase), in agreement with *in silico* DegScore predictions. Together, these results constitute a proof of concept that reinforcement learning is a viable, training-data-free framework for rational mRNA vaccine design.

## Introduction

mRNA vaccines and mRNA-based therapeutics have emerged as versatile platforms for a wide range of tasks such as infectious disease targeting, cancer immunotherapy and protein replacement therapy (Chaudhary et al., 2021; Malonis et al., 2020; Parhiz et al., 2024; Xia, 2021). Their efficacy is governed not only by the choice of target but also by intrinsic properties of the manufactured RNA molecule (Jin et al., 2025; Leppek et al., 2022; Zhang et al., 2023, 2025). For example, codon usage bias in mRNAs reflects the non-uniform use of synonymous codons driven by the heterogeneous, cell-type-dependent tRNA pool. In line with this, the <u>C</u>odon <u>A</u>daptation Index (CAI) is the most widely adopted metric of codon suitability for mRNA vaccine design used as a proxy for optimizing cellular translation efficiency (Bahiri-Elitzur & Tuller, 2021; Hanson & Coller, 2018; Roth, 2012; Sharp & Li, 1987). Similarly, the thermodynamic stability of the mRNA influences molecular half-life and the kinetics of protein production (more stable structures support long-term output, whereas less stable structures favor a rapid initial burst of translation), and can be estimated from its <u>M</u>inimum <u>F</u>ree <u>E</u>nergy (MFE) (Juszkiewicz et al., 2020; Leppek et al., 2022; Mauger et al., 2019).

Since intracellular stability of mRNA directly affects the total amount of protein that can be generated from it and fast degradation remains a practical bottleneck for storage and distribution of RNA-based therapies, improving codon usage (via CAI) and thermodynamic stability (via MFE) through sequence design is an alternative strategy for obtaining more efficient, stable, and deliverable mRNA vaccines (Cheng et al., 2023; Kim et al., 2024; Pfeifer et al., 2023; Romero et al., 2016). However, simultaneous optimization of both features is a combinatorial search problem with a large solution space because a single protein can be encoded by an enormous number of synonymous mRNAs due to the codon degeneracy. For example, the SARS-CoV-2 spike protein can be coded by more than 10⁶³² synonymous mRNAs (Kim et al., 2024). Crucially, CAI and MFE are in tension: maximizing codon bias can erode intramolecular base pairing, while stabilizing the RNA structure often requires rarer codons (Kim et al., 2024; Zhang et al., 2023, 2025). Thus, RNA therapy design is a multi-objective optimization problem in which one looks for a solution on the Pareto front that, while not reaching the maximum of either objective, achieves an efficient compromise between them. Moreover, the combinatorial explosion of possibilities renders brute- force search infeasible. This explosion in possible RNA designs worsens in the case of multi- epitope vaccines, where the spatial order of epitopes within a single construct creates multiple additional, non-equivalent solution spaces (Jin et al., 2025; Shahraki et al., 2025).

Several computational tools have tried to address this problem. Codon optimization tools target translational efficiency, increasingly using machine learning (Fu et al., 2020; Jain et al., 2023), while structure-focused tools optimize the stability of the mRNA’s secondary structure (Krueger & Ward, 2024; Leppek et al., 2022; Terai et al., 2016). Few tools optimize both parameters simultaneously. A remarkable recent case is the deterministic pipeline LinearDesign, which introduced the MFECAIλ objective function, a score that balances MFE and CAI through a tunable λ parameter (Zhang et al., 2023). Also, some new pipelines make use of deep generative approaches (Zhang et al., 2025). These new methods have demonstrated that MFE-prioritized designs improve stability and immunogenicity. However, deterministic single-pass methods can only return one candidate per input, leaving unexplored near-optimal synonymous solutions that can only be discovered by stochastic approaches.

We tackled this problem developing ReinforSeqPipe, a computational pipeline that exploits <u>R</u>einforcement <u>L</u>earning (RL) to optimize multiepitope cancer mRNA vaccine sequences. Our results suggest that RL-based methods improve the MFECAIλ significantly over codon-optimized controls, and that parallelization allows ReinforSeqPipe to outperform the state-of-the-art of deterministic algorithms.

## Materials and Methods

### ReinforSeqPipe workflow

ReinforSeqPipe optimizes mRNA sequences with a deep Q-network (DQN) reinforcement learning agent. The agent uses a bi-parametric minimum free energy–codon adaptation index (MFECAIλ) score as its reward function (Mnih et al., 2015, Figure 1). The agent requires no training dataset; it learns solely from reward signals. The pipeline comprises numerical encoding of the input sequence, a <u>L</u>ong <u>S</u>hort-<u>T</u>erm <u>M</u>emory (LSTM)-based state encoder, a DQN agent with ε-greedy exploration, and *in-silico* evaluation through the reward function (Figure 1).

**Figure 1.**
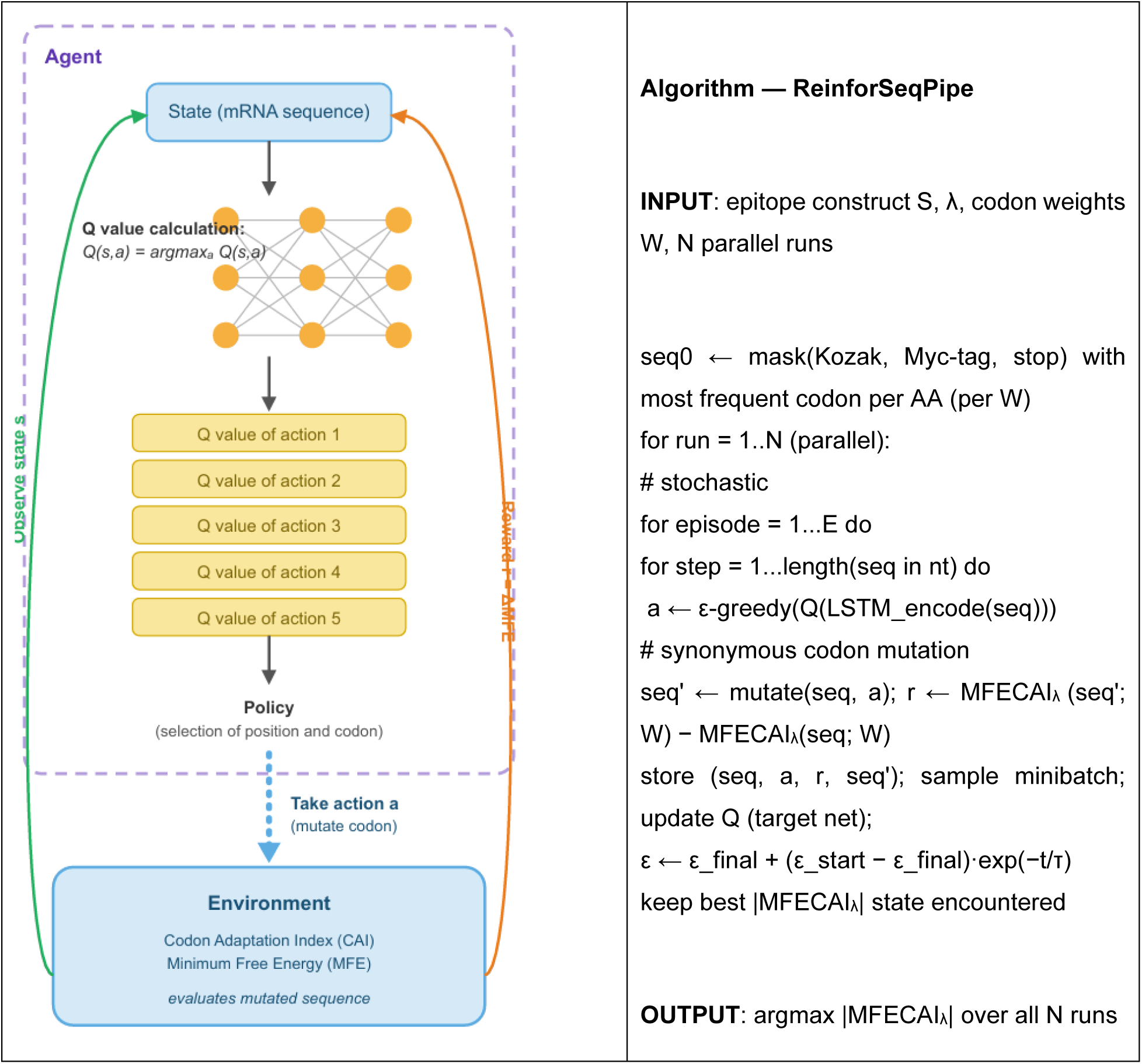
**Left. Deep Q-Network (DQN) architecture of the ReinforSeqPipe pipeline**. The reinforcement learning loop treats the current mRNA sequence as the state (s). A deep neural network estimates Q-values for candidate synonymous mutations, and the policy selects the next action (a)—mutating a specific codon at a specific position—and applies it to the mRNA. The environment evaluates the mutated sequence via its codon adaptation index (CAI) and minimum free energy (MFE), returning the reward r = ΔMFECAIλ, MFECAI*_λ_* = MFE − *λ* ⋅ *L* ⋅ log(CAI). The agent then observes the new state and the loop repeats. **Right. Pseudocode of the ReinforSeqPipe pipeline**.

### Sequence encoding

We one-hot encoded each codon as a 12-dimensional vector (four possible bases per nucleotide × three nucleotides) and stacked the vectors into a 12 × N matrix, where N is the sequence length in number of codons. An LSTM encoder processed the sequence- encoding matrix to capture long-range dependencies between distant codons. This property makes the LSTM a better alternative to conventional recurrent neural networks on long sequences, owing to its higher resistance to gradient problems during backpropagation (Van Houdt et al., 2020). We fed the learned state representation into the DQN agent to evaluate the effect of each mutation on the state and reward.

### Codon weights

We retrieved raw codon counts from the TissueCoCoPUTs database (Kames et al., 2020), and we computed the weight of codon c as

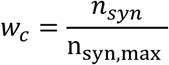

Where *n_syn_* is the occurrence count of codon *c* and n_syn,max_ is the count of the most frequent synonymous codon. We used the "Genomic" weight dictionary as the default for CAI calculation because it derives from average gene expression across all human tissues. We employed tissue- specific dictionaries ("Whole Blood", "Skin sun exposed", "Skin not sun exposed") to test sensitivity. We derived the codon usage tables from the CoCoPUTs database, specifically the TissueCoCoPUTs database (Kames et al., 2020).

### Biophysical metrics

We computed MFE and secondary structure with ViennaRNA v2.7.0 as a Python Library (Lorenz et al., 2011), and computed CAI as

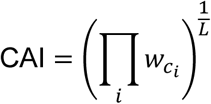

with *L* is the length of the mRNA in codons (Sharp & Li, 1987). We computed GC content, GC3 (Shen et al., 2015) and the effective number of codons (ENC) (Wright, 1990) with the Codon Bias Python library (Diament, 2022). ENC was calculated from the average homozygosity using the formula

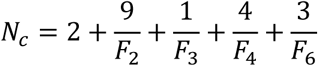

where F2–F6 represent the average homozygosity for each degeneracy. These classes measure the mean codon usage homozygosity for amino acids grouped by their degree of redundancy (degeneracy) in the genetic code.

### Reward function

We used the reward defined in the LinearDesign pipeline (Zhang et al., 2023):

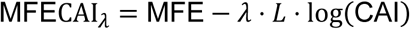

where λ balances the two objectives: λ = 0 prioritizes MFE, while λ >> 1 prioritizes CAI. *L* is the length of the RNA in codons. To support interpretation of the model and comparison of results, we used the absolute value of the MFECAIλ score:

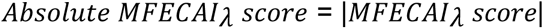

The agent’s task was to maximize this absolute value through synonymous mutagenesis.

### DQN and exploration

The DQN implements experience replay and a target network that we update less frequently than the primary network to stabilize learning. It uses the standard temporal-difference loss (Mnih et al., 2015; Tsitsiklis & Roy, 1997). We selected actions with an ε-greedy policy with exponential decay (Boyan & Moore, 1995) ,

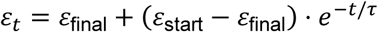

where τ is the decay factor multiplied by the steps per episode. Because ε-sampling and replay batching are stochastic (Nagarajan et al., 2019), we optimized each sequence in triplicate and retained the best of the three solutions. The implementation utilized built-in PyTorch classes.

### Hyperparameter optimization

We derived the values above through a two-stage procedure on a representative construct (see below). First, we performed a one-at-a-time sensitivity screen of eight hyperparameters (defaults: decay rate 1.5, update rate 10, hidden size 128, batch size 64, start ε 1.0, final ε 0.05, learning rate 0.001, discount factor 0.95; 50 runs per setting). We carried forward the hyperparameters that showed a significant effect (one-way ANOVA); we did not test the others further. Second, a grid search over 192 combinations of learning rate, hidden size, batch size, and update rate (n = 50 each) identified the configuration with the best mean MFECAIλ at acceptable runtime: learning rate 0.01, hidden size 128, batch size 64, update rate 10 (∼232 s per optimization on an Nvidia RTX A6000, see Sup. Table S1). Notably, neither the ε-greedy parameters nor the discount factor significantly affected performance, which suggests that the optimization environment is readily learnable and that the Q-values stabilize early. Representative construct: YLEPGPVTAGGGGSEVDPIGHLYSLYNTVATLILKEPVHGVGGGGSEQKLISEEDL

### Computational setup and runtime analysis

We performed all computations on a workstation with an NVIDIA Quadro RTX A6000 GPU (48 GB), using PyTorch for the DQN/LSTM components and ViennaRNA v2.7.0 for structure and MFE calculation. We profiled runtime on 17 mRNAs of increasing length (59–211 amino acids; Sup. Table S2). Total time grew approximately exponentially with length, and ViennaRNA calls dominated the budget, because each step requires a full folding computation. Single optimizations stay within one hour for mRNAs up to ∼750 nucleotides. LinearDesign optimizes comparable sequences in seconds; this trade-off is offset by ReinforSeqPipe’s parallelizable best-of-N sampling.

### Construct design

Constructs comprised 1) a Kozak consensus sequence with AUG start codon (Kozak, 1991), 2) five established HLA-A*02:01-binding peptides (originating from proteins GP100, MelanA, MageA3, HIV-gag, HIV-pol) joined by GGGGS linkers, 3) a constant in-frame Myc-tag (EQKLISEEDL) for detection at the C-terminal, 4) a UAA stop codon chosen for its superior termination efficiency (Lee et al., 2021), and 5) a poly A tail of a defined length of 64 A. The Myc-tag and stop codon were masked from mutation. We optimized all 120 (5!) permutations of peptide order. Initial sequences used the most frequent codon per amino acid from the "Genomic" table due to universality. Each episode comprised 249 steps (sequence length in nucleotides) over 10 training episodes.

### λ tests and benchmarking

We benchmarked ReinforSeqPipe at λ ∈ {1, 4, 10}, as in (Zhang et al., 2023). To ensure fairness, benchmarks against LinearDesign used the pipeline’s native codon weights. Being deterministic, LinearDesign ran once per sequence, while ReinforSeqPipe (stochastic) ran up to 50 times. For the benchmarking set, each optimization had 20 episodes of training. Dedicated convergence experiments motivated this 20-episode budget: performance improved with episode count up to 20, showed only a small gain between 20 and 25 episodes, and showed no significant improvement at 30 episodes with linearly increasing runtime. Correspondingly, a stopping criterion that halts training after five consecutive episodes without improvement typically terminated runs by episode 25.

### Experimental validation

Plasmid constructs were ordered from Thermofisher’s Geneart division and transformed into *E. coli* XL1-Blue (Sigma-Aldrich, United States). After replication in bacterial culture followed by isolation, they were transcribed *in vitro* with the mMESSAGE mMACHINE T7 ULTRA kit (Thermo Fisher Scientific, United States) according to the manufacturer’s instructions. The resulting mRNA was purified with the Mini RNeasy kit (Qiagen, Germany) and quantified it on a NanoDrop 2000c (Thermo Fisher Scientific, United States). We performed electroporation on a GenePulser XceII (Bio-Rad, United States) as described previously (Sauerer et al., 2024). We quantified mRNA decay by RT-qPCR at 0.5, 1-, 2-, 4-, 6-, and 24-hours post-electroporation, normalized to GAPDH (n = 4 biological replicates). We ran reactions using the innuMIX Standard PCR Master Mix (Analytik Jena, Germany) on a qTower3 thermal cycler (Analytik Jena, Germany). We generated cDNA with the SuperScript IV VIVO Master Mix (Thermo Fisher Scientific, United States).

## Results

### ReinforSeqPipe, a Deep Q-Network (DQN)-based reinforcement learning pipeline to design RNA therapeutics

We present ReinforSeqPipe, a DQN-based reinforcement learning pipeline that simultaneous optimizes CAI and MFE for mRNA vaccine sequences (**Figure 1-2**). Reinforcement learning (RL) offers a promising framework for optimizing RNA sequences: an agent improves a policy purely from the reward signal it receives from the environment, so it requires no labeled training data (Mahmud et al., 2018). DQNs, originally developed for video games AI, perform well in high-dimensional action spaces and have been applied to biology- related problems (Mahmud et al., 2018; Mnih et al., 2015). In our approach, we formulate the sequence design task as a Markov decision process (Amodu et al., 2025), using the standard RL decomposition (**Figure 1**). The mRNA sequence defines the **state** (*s*), with MFE and CAI as state features. Synonymous mutations constitute the **actions** (*a*) applied to the state. The change in the MFECAIλ score after each mutation defines the **reward** (*r*). The agent samples actions according to its policy, receives reward signals after each step, and updates Q-values to maximize cumulative returns. Through this iterative interaction, ReinforSeqPipe discovers beneficial mutations directly from the biophysical objective, without any supervised training data.

**Figure 2.**
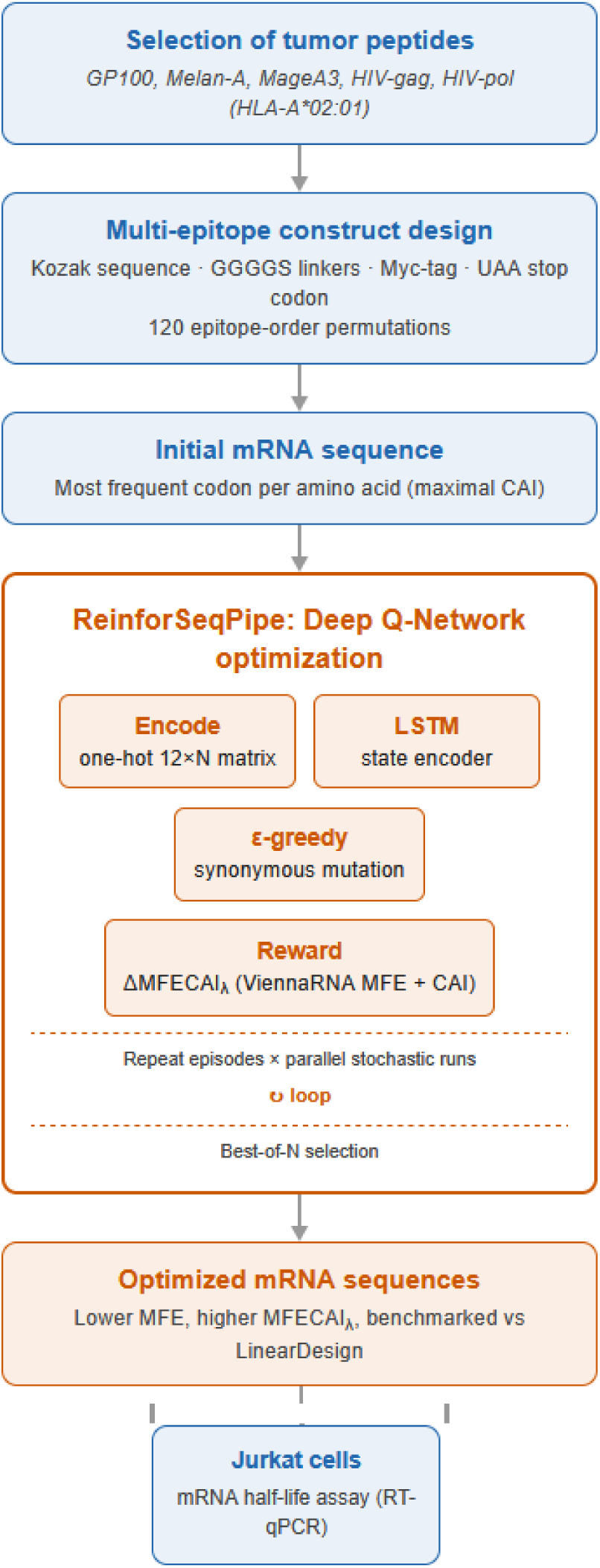
**Overview of the ReinforSeqPipe workflow**. First, the peptides to integrate into the RNA sequence are selected by manual curation, experimentation, or computational algorithms (Lischer et al., 2024). To establish the workflow, we here chose 5 model peptides from cancer and viral antigens. We then construct the multi-epitope by joining peptides with GGGGS linkers, flanking it with a Kozak consensus sequence and an in-frame Myc-tag followed by a UAA stop codon, and generate all 120 epitope-order permutations. The pipeline builds initial mRNA sequences from the most frequent codon per amino acid and optimizes them through synonymous mutagenesis with a DQN agent. Stochastic runs execute in parallel and we retain the best-of-N solution and benchmark it against LinearDesign. We validate optimized constructs by measuring mRNA half-life in Jurkat cells (RT-qPCR).

### ReinforSeqPipe improves multi-epitope mRNA sequence parameters

We optimized all 120 permutations of a five-epitope vaccine construct (**Figure 2).** Because ReinforSeqPipe is stochastic, we optimized each sequence three times and retained the best resulting variant. Optimized sequences achieved a mean MFECAIλ of 145.6 ± 5.8 versus 109 ± 1.8 for the unoptimized controls (paired t-test, p < 0.001; **Figure 3**D). The improvement came mainly from a significant increase in MFE (data not shown), while CAI decreases modestly. Previous studies discussed this MFE–CAI trade-off (Kim et al., 2024; Zhang et al., 2023). The optimized scores shifted upward and spread much wider than the control scores (SD 5.8 vs 1.8). Two phenomena likely explain this larger spread: the expected large number of near-optimal synonymous solutions, and the partially distinct solution space of each peptide permutation. Final and initial scores did not correlate (Pearson r = 0.08, p = 0.38; Sup. Mat. Figure S1), so ReinforSeqPipe did more than refine the initial sequences.

**Figure 3.**
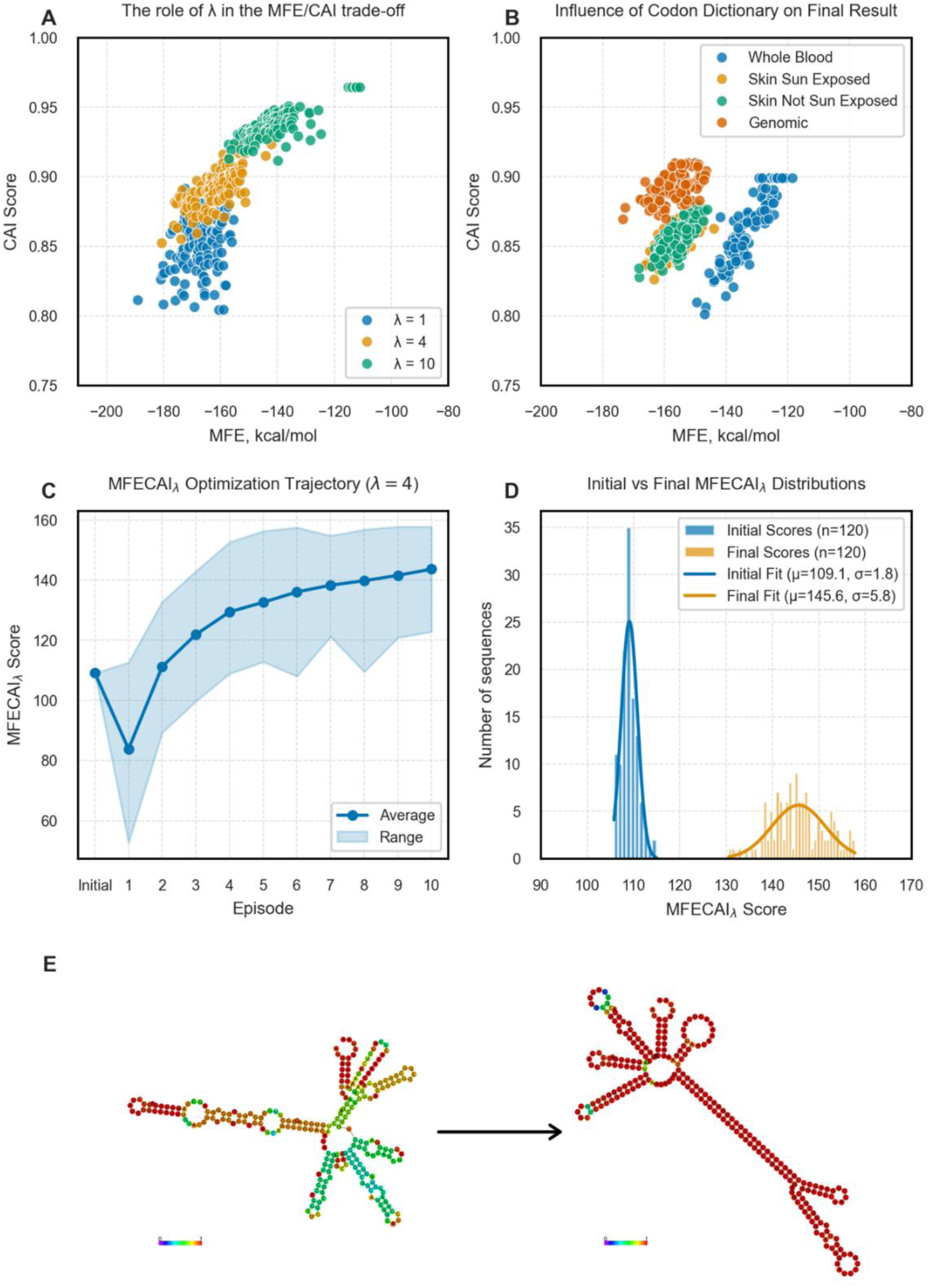
A. The joint scattering plot of best of three datasets. Each dot represents the best of three optimized variants for each of its permutation**. B. The role of Relative Codon Usage in the performance and final solutions.** The initial sequences were also assembled according to the relevant Codon Usage. **C. Optimization progresses across episodes of training**. In total 10 episodes were used to find the best solution. The range shows the range of parameters for all of 120 permutations during the optimization process. **D. The distributions of MFECAIλ scores for Initial (control) and Final (optimized) groups** (best-of-three set). Additionally, mean and standard deviation values are shown. **E.** ViennaRNA-predicted secondary structure showing the initial state and the best optimized variant based on MFECAIλ.

For codon-related parameters CAI, GC3, ENC, and GC, testing these metrics was impractical because the control group had zero variance in codon composition. However, the general trends point to successful optimization. Final solutions showed higher codon diversity (indicated by increased ENC), consistent with codon substitutions that reconfigure hybridization and strengthen thermodynamic stability. GC and GC3 changes were smaller, but the final solutions are described as GC-rich (GC ≥ 70%, GC3 ≥ 80%).

Across ten episodes, the learning and optimization time course showed the expected exploration– exploitation pattern. MFECAIλ dropped sharply at episode 1 (**Figure 3C)** because of losses in both codon optimality and thermodynamic stability (data not shown), as the agent explored the environment without prior knowledge. Next, the objective function rose gradually and plateaued near episode 10. Sustained score gains came mainly from the MFE trajectory, while CAI stabilized early with diminishing marginal losses. The plateau near episode 10 indicates that the agent had largely converged to a stable mutation strategy, with marginal gains diminishing over the final episodes.

We selected the three post-optimization sequences with the highest MFECAIλ, together with their synonymous controls, for closer inspection. The optimized sequences showed much lower predicted MFE than their controls (-175.4 to -180.6 kcal/mol and -111.5 to -117.1 kcal/mol, respectively, **Figure 3D**). CAI decreased from 0.964 to 0.882, 0.852, 0.880 respectively (See Sup. Mat Table S3). Predicted 2D structures showed the visual improvement of the base-pairing probability for the best optimized variant compared to its initial state (**Figure 3E**). The second best and the third best mRNAs also showed an improvement (See Sup. Mat Figure S3 and Figure S4).

### The weighting factor λ defines the balance between structural and codon optimality

To evaluate the effect of modulating the importance of each subfunction in the results (CAI- and MFE-centered ones), we optimized all sequences at three different λ values (1, 4, and 10), following the original formulation in the LinearDesign pipeline (Zhang et al., 2023). We ran each optimization in triplicate. The joint distribution of CAI and MFE across the 120 best-of-three solutions followed the trajectory the objective function predicts (**Figure 3**): setting λ to 1 produced solutions with the most negative MFE (prioritizing thermodynamic stability), setting it to 10 resulted in solutions with the highest CAI (prioritizing codon adaptation), and a value of λ equal 4 favored a compromise between the MFE and CAI.

When favoring MFE optimality, the distributions of both CAI and MFE occupied comparatively wider intervals. In other words, when structural stability dominates, the agent appears less constrained in incorporating rarer codons, thereby increasing codon diversity and the hybridization space of feasible structures. Interestingly, tuning a single parameter, λ, consistently shifted all 120 optimized sequences along the predicted trajectory. This finding supports usage of λ as a practical design dial: decrease λ for thermodynamically stable constructs suited to moderate long-term expression or increase λ for CAI-oriented constructs suited to burst translation (**Figure 3A**).

### Codon weights reshape the accessible solution space

To probe the influence of the codon reference on the outcome, we optimized the 120-permutation dataset under four codon-weight dictionaries derived from the TissueCoCoPUTs database ("Genomic", "Whole Blood", "Skin sun exposed", and "Skin not sun exposed"), with initial sequences assembled from the respective tables. The choice of dictionary shifted both the range and the dispersion of the final solutions in MFE–CAI space, with the "Whole Blood" weights producing markedly wider distributions than the other three (**Figure 3B).** This suggests that codon weights do more than rescale the CAI term: they alter the hybridization landscape accessible to a given polypeptide, effectively relocating the boundary of the feasible region in the MFE–CAI plane. Because of its tissue-universality, we retained the "Genomic" dictionary for all subsequent optimizations and for the experimentally validated constructs.

### Benchmarking against LinearDesign

To evaluate whether ReinforSeqPipe is competitive with state-of-the-art tools, we benchmarked it against the deterministic algorithm LinearDesign (Zhang et al., 2023). The benchmark used the same objective function and 120-permutation sequence dataset, with codon weights taken from the LinearDesign pipeline to avoid any related bias. In the design, the Myc-tag is fixed at the C-terminus. We built the initial ReinforSeqPipe sequences from the most frequent codon per amino acid, giving an initial CAI of 1. Because LinearDesign is deterministic, we ran it once per sequence. However, ReinforSeqPipe is stochastic, and therefore we ran it up to 50 times per sequence and tracked the number of iterations needed to beat the LinearDesign solution. This asymmetric approach tests whether the best-of-N strategy available to a stochastic, parallelizable algorithm compensates for the replicability of a deterministic one. We report results for the three λ settings: MFE-biased (λ=1), Balanced (λ=4), and CAI-biased (λ=10).

At MFE-biased setting λ=1, the solutions clustered in approximately the same MFE interval (**Figure 4A**). Nevertheless, a single ReinforSeqPipe run outperformed LinearDesign in 84 of 120 permutations. By 10 repeats, it outperformed LinearDesign in all 120 (**Figure 4B**). Given the similar location of solutions in MFE-CAI space, the results suggest that ReinforSeqPipe finds more stable variants.

**Figure 4.**
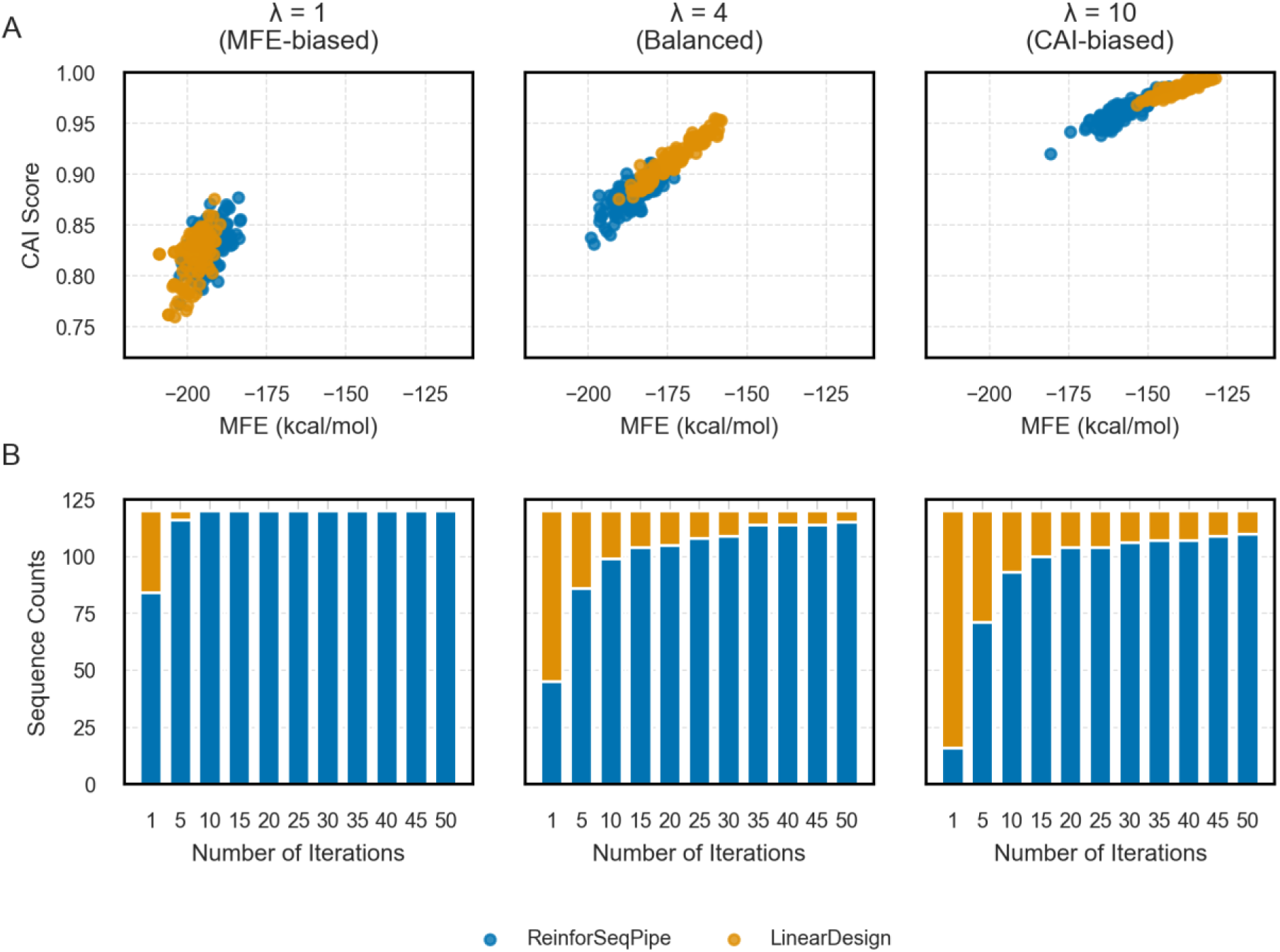
A. Scatter plots comparing the positions of solutions in MFE–CAI space, showing the difference between the ReinforSeqPipe and LinearDesign pipelines in codon adaptation and secondary-structure stability. We selected the best of 50 ReinforSeqPipe solutions. **B. Barplot comparison** of ReinforSeqPipe and LinearDesign in this work’s setup. Each bar shows which algorithm performs better for each of the 120 permutations.

At the Balanced setting λ=4, comparative scatter plots in MFE–CAI space showed a consistent pattern: ReinforSeqPipe solutions sit at slightly better MFE with marginal CAI loss compared to LinearDesign (**Figure 4A**). LinearDesign remained dominant at a single repeat, but ReinforSeqPipe took the lead by 5 repeats (86/120) and led 115/120 at 50 repeats (**Figure 4B**).

Similar to the Balanced setting, at the CAI-biased setting λ=10 ReinforSeqPipe lagged behind LinearDesign at a single repeat, showed no significant difference at 5 repeats, led at 10 repeats (93/120), and remained dominant in 110/120 at 50 repeats **(Figure 4B).** Since a high λ value correlates with greater codon variability, our results suggest that ReinforSeqPipe performs better when the search space expands via codon variability.

Practically, a modest number of parallel runs (≤ 10) is enough for ReinforSeqPipe to match or beat most deterministic solutions at every λ setting, and the advantage is largest where structural optimization matters most. Moreover, ReinforSeqPipe seems to be learning and implementing a strategy that favors marginal stability gains over codon-adaptation gains when the two trade off.

### Compute cost

To characterize scalability, we profiled computation time on 17 mRNAs of increasing length (59–211 amino acids; UniProt entries listed in Sup.Mat. Table S2). Total runtime grew steeply — approximately exponentially — with sequence length. Timing of individual ViennaRNA calls showed that structure and MFE evaluation dominate the cost, while the remaining RL components (Q-value estimation, reward calculation, replay and encoding) contribute a smaller fraction. According to the exponential fit, a single optimization stays below one hour for mRNAs of up to ∼750 nucleotides. LinearDesign, by contrast, optimizes a single sequence of this size in seconds; its advantage in wall-clock time is clear. We consider this an acceptable trade-off, since both tools target vaccine-design decisions where minutes-to-hours of computation are negligible relative to downstream experimental costs.

### Experimental validation

To test whether an increase in predicted thermodynamic stability increases RNA half-life in living cells, we compared the best optimized sequence against two controls: its synonymous mRNA, and an mRNA coding for another permutation variant built from the same codon set. The relative mRNA levels, normalized by GAPDH, were plotted against time course and their half-lives were calculated (4 biological replicates per sequence; **Figure 5)**. We then calculated individual half-lives from the abundance time courses. The decay curve of the optimized mRNA remained above both controls throughout the 24 hours period, and its derived half-life was significantly longer than those of the synonymous control (Initial 1) and of the non- synonymous control (Initial 2) (Mann-Whitney U-test; **Figure 5A**).

**Figure 5.**
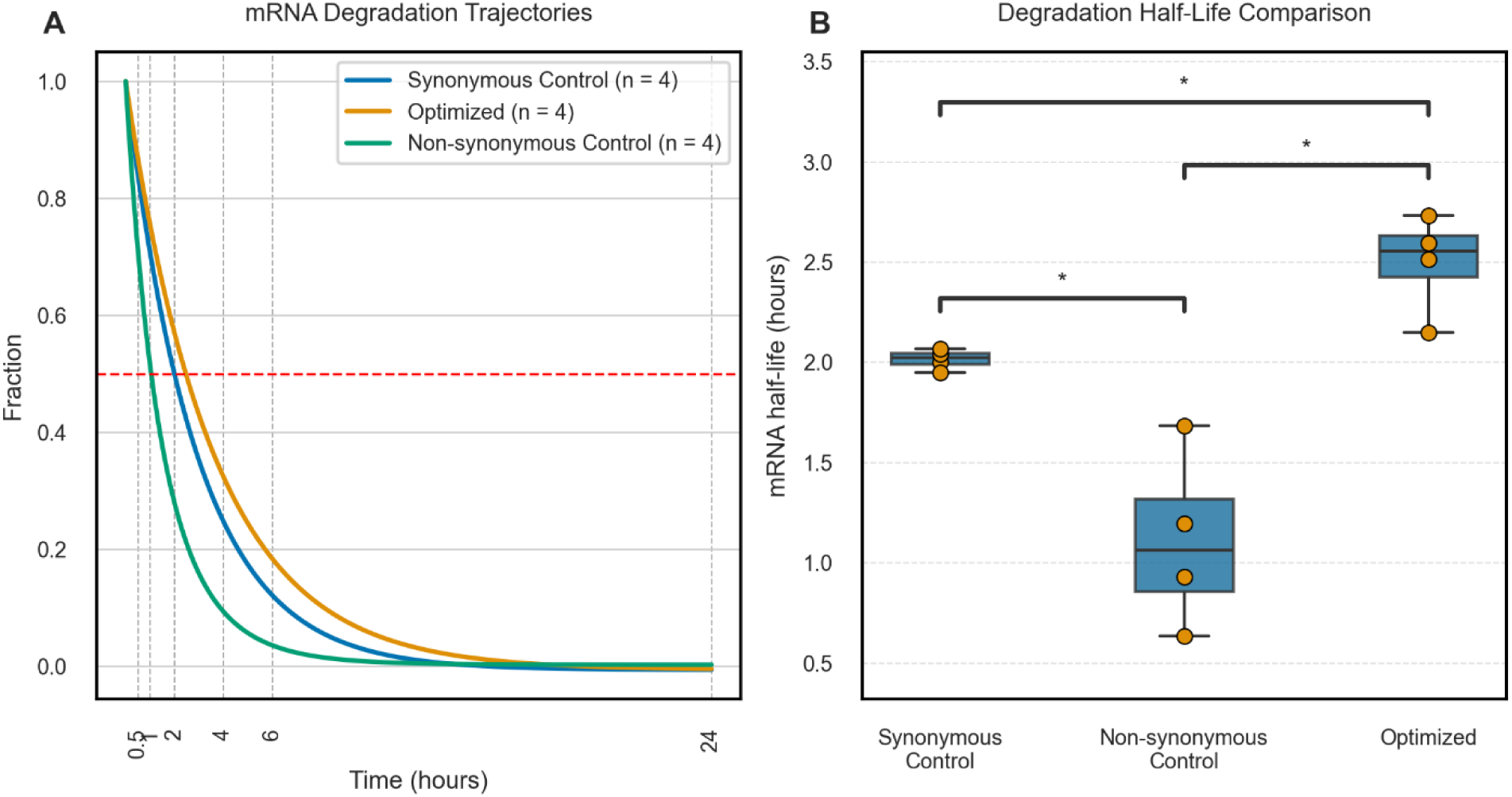
A. Projected decay curves of the tested mRNAs based on qPCR results. The decay curves are based on the fraction of remaining mRNA at specific time points (grey vertical dash lines) against GAPDH levels: 0.5-, 1-, 2-, 4-, 6-, and 24-hours. The red horizontal dash line shows the half fraction of original amount of mRNA**. B. The mRNAs half-lives comparison**. The graph shows the half-life of three different mRNA sequences. The Synonymous control and Optimized mRNAs share the same polypeptide variant, meanwhile the non-Synonymous control codes for the different polypeptide. Each experiment was repeated 4 times.

Two findings from this experiment require discussion. First, the non-synonymous control degraded substantially faster (∼1 h shorter half-life) despite having a predicted MFE very similar to the synonymous control. Global MFE therefore does not fully determine cellular mRNA stability, a feature that is consistent with reports that local structural motifs can drive accelerated degradation (Wayment-Steele et al., 2022). Second, the wide spread between replicates (**Figure 5B**) likely reflects the small replicate number (n = 4) and technical variance from the multi-step protocol using living cells (noise arising from electroporation efficiency, metabolic activity of the cells, efficiency of RNA extraction and purification, reverse-transcription efficiency, and qPCR primer binding). The optimized and synonymous constructs encode the same polypeptide, so the half-life difference is attributable to the sequence design. As an orthogonal, *in silico* check, the DegScore pipeline (Leppek et al., 2022) predicted the optimized constructs to have, on average, half-lives 12-minute longer and lower degradation scores than their controls (Supplementary Figure S4), consistent with our experimental measurements.

## Discussion

ReinforSeqPipe shows that deep reinforcement learning offers a viable, training-data-free framework for simultaneous optimization of codon adaptation and thermodynamic stability in mRNA vaccine design. The method frames synonymous sequence design as a Markov decision process: the sequence is the state, a synonymous mutation is the action, and the change in the MFECAIλ score is the reward. The agent then learns beneficial mutational strategies directly from the biophysical objective, with no supervised dataset. Across the 120 epitope-order permutations of a five-epitope construct intended for HLA-A*02:01 presentation, the pipeline raised MFECAIλ substantially over codon-optimized controls and modulated the MFE–CAI trade-off in a manner consistent with theoretical predictions. It also matched or exceeded the deterministic state-of-the- art LinearDesign algorithm at all tested λ values within ten stochastic repeats. Most importantly, the predicted stability gains produced a measurably longer cellular mRNA half-life in Jurkat cells.

### Relation to existing methods

Classical codon optimization tools maximize translation-related scores such as CAI and, at best, penalize sequence features known to impair expression; however, they largely ignore RNA structure. Structure-oriented pipelines take the opposite route. LinearDesign remains the strongest recent reference point for our algorithm (Zhang et al. 2023). LinearDesign introduced the MFECAIλ objective that we adopt unchanged. The basis of its algorithm is to perform dynamic programming over a codon graph and find, for a given λ, the probably optimal sequence under its own scoring scheme.

On a per-run basis, a deterministic global optimizer such as LinearDesign retains a clear advantage in terms of computation time. ReinforSeqPipe’s strength lies elsewhere. First, ReinforSeqPipe samples the solution space: each stochastic run yields a different near-optimal candidate. Our parallel execution therefore implements a best-of-N strategy that, in our results, recovered or exceeded LinearDesign’s solution on the majority of permutations at every λ tested. This feature can be a practical advantage for ReinforSeqPipe: in some scenarios, vaccine development benefits from diverse candidate sequences—for instance, to navigate patent constraints, manufacturing considerations, or pre-existing immunity against encoded epitope junctions. Moreover, ReinforSeqPipe is deliberately modular: it establishes the design principle that stochastic, reward-driven sequence exploration can productively navigate the vast synonymous search space. Second, the sampling behaviour is adaptive: at λ = 1 (MFE-centric), ReinforSeqPipe found solutions with comparable MFE but better CAI than LinearDesign. This suggests the agent implicitly learns to escape the codon homogeneity that pure MFE maximization imposes. Third, MFE evaluation of candidate mutations dominates the per-step cost here. Our analysis confirms that ViennaRNA calls constitute the runtime bottleneck and that total time grows steeply with sequence length. An accelerated or approximate RNA folding predictor therefore offers the most direct route to scaling and accelerating the pipeline.

More recent deep generative approaches likewise operate without exhaustive search, but typically require large corpora for pretraining, which biases them towards naturally observed coding sequences. ReinforSeqPipe sits at an attractive middle point: no training data, an interpretable, physically grounded reward, and a hyperparameter budget small enough (hidden size 128, ∼250- step episodes) to run on standard, local hardware.

Additionally, ReinforSeqPipe constitutes an example of how Reinforcement Learning can mitigate the problems of imbalanced or incomplete biological datasets with a good reward function.

### The λ parameter as design dial controlling RNA specifications

Low λ values push solutions towards thermodynamically stable constructs suited to sustained, long-term RNA output. In contrast, high values favour codon-adapted constructs for rapid translational bursts. MFE- prioritized optimization produced broader CAI and MFE distributions. This fits the intuition that structural optimization is the harder sub-problem: when stability dominates, the agent freely incorporates rarer codons, which enlarges both codon diversity (higher ENC) and the accessible hybridization space. We note, however, that all optimized solutions remained GC-rich, although this study did not evaluate GC-driven effects on innate immune sensing and translation.

### Experimental validation

The optimized mRNA decayed significantly slower than both a synonymous control and a non-synonymous control encoding a different epitope permutation from the same codon pool. This translates the ∼60 kcal/mol predicted MFE advantage into a roughly 20-minute half-life extension in Jurkat cells. Two observations deserve emphasis. First, the non- synonymous control degraded faster despite near-identical predicted MFE to the synonymous control. This is aligned with reports that global MFE does not fully determine cellular stability and that local structural motifs—ribosome pause sites, endonuclease-accessible regions—can dominate degradation kinetics. Second, the small replicate number and the multi-step protocol followed (electroporation efficiency, column purification, RT and primer efficiencies) contributed substantial variance: the stability claims, while statistically supported, require confirmation with more replicates and direct RNA quantification (e.g., branched-DNA assays avoiding amplification bias). The data also need to be extended to protein-output measurements to confirm the relevance of the CAI score, which link stability to the quantity that actually matters therapeutically, and ultimately to T cell stimulation.

### Limitations

The current version of the pipeline optimizes based on sequence-level proxies for RNA features (MFE from unmodified nucleotides and CAI from bulk tissue codon tables). Clinically deployed mRNAs carry heavy nucleoside modifications (e.g., N¹-methylpseudouridine), are divided into UTRs and a coding region, and include cap and poly(A) structures, all of which influence RNA stability (Wu et al., 2026). Computational secondary-structure prediction carries some intrinsic uncertainty, particularly at the GC levels we reach here (Zuber & Mathews, 2019). In our benchmark, we used an asymmetric protocol (one LinearDesign run vs. up to fifty stochastic ReinforSeqPipe runs). Although the deterministic/stochastic distinction justifies this asymmetry, we concede that it may inflate LinearDesign’s apparent advantage in wall-clock-comparable terms.

### Outlook

Three directions seem most promising for improving the algorithm. First, one can replace the single MFE evaluator with richer reward components—e.g., ensemble defect, uORF avoidance, and GC windows (Imdad et al., 2023; Jin et al., 2025). This can convert ReinforSeqPipe’s reward from bi-parametric to genuinely multi-objective. In that case, one would have to compute the Pareto front of the problem, with λ as a vector instead of a scalar, and the agent as a sampler of the frontier. Second, more advanced RNA folding models, including learned structure predictors and dynamic folding algorithms, can be integrated as evaluators (Zhao et al., 2010). However, the computational time of these methods is anticipated to be the main bottleneck of future version of ReinforSeqPipe. The CAI subfunction currently draws its codon weights from generic, bulk-tissue frequency tables, whereas optimal codon usage differs between tissues. One could use inverse reinforcement learning to infer tissue-appropriate weights from the genes most highly expressed in the target tissue. This way, the reward function would become tissue-specific and data-driven rather than static. Finally, from an experimental perspective, one could improve the validation measuring protein expression output and T-cell activation responses. These additional experiments will determine whether the computational gains we report here translate into therapeutic benefit.

## Data and code availability

The codon-weight tables (Genomic, Whole Blood, Skin sun exposed, Skin not sun exposed, LinearDesign), the optimized and control sequences, and the ReinforSeqPipe source are available at the GitHub repository https://github.com/JVeraLab/reinforseqpipe.

## Contributions

JV and JD developed the concept. BM and JV sketched the computational approach. BM designed, implemented, trained and validated the computational model under the supervision of JV. BM performed the experiments under the supervision of JV and JD. BM, ME, and JV drafted the manuscript. JV and CB acquired the funds. All authors edited and approved the submitted draft.

## Ethics approval and consent to participate

The studies involving human subjects or material were reviewed and approved by the Ethics Committee of the Friedrich-Alexander Universität Erlangen-Nürnberg (ethics approval no. 4158). The principal investigator and all other investigators ensured that the studies were conducted in full conformity with the principles set forth in the Declaration of Helsinki.

## Funding

Open Access funding enabled and organized by Project DEAL. JV was supported by the Forschungsstiftung Medizin Uniklinikum Erlangen, the German Federal Ministry of Research, Technology and Space (BMFTR) via projects e:Med MelAutim [01ZX1905A, 01ZX2205A] and KI- VesD [161L0244A, 16LW0338K], and the Bavarian Cancer Research Center (BZKF) through the project PROMIT [BGF/25/06/LMU/Heinz]. JV was supported by the Hiege-Stiftung – Die Deutsche Hautkrebsstiftung through the project "Melanoma risk classification based on multi-level clinical and molecular profiling data and artificial intelligence".

## Acknowledgements

We thank the support of Niels Schaft and Annett Hamann. Lumo AI was used to proofread the manuscript.

---

**Figure S1.**
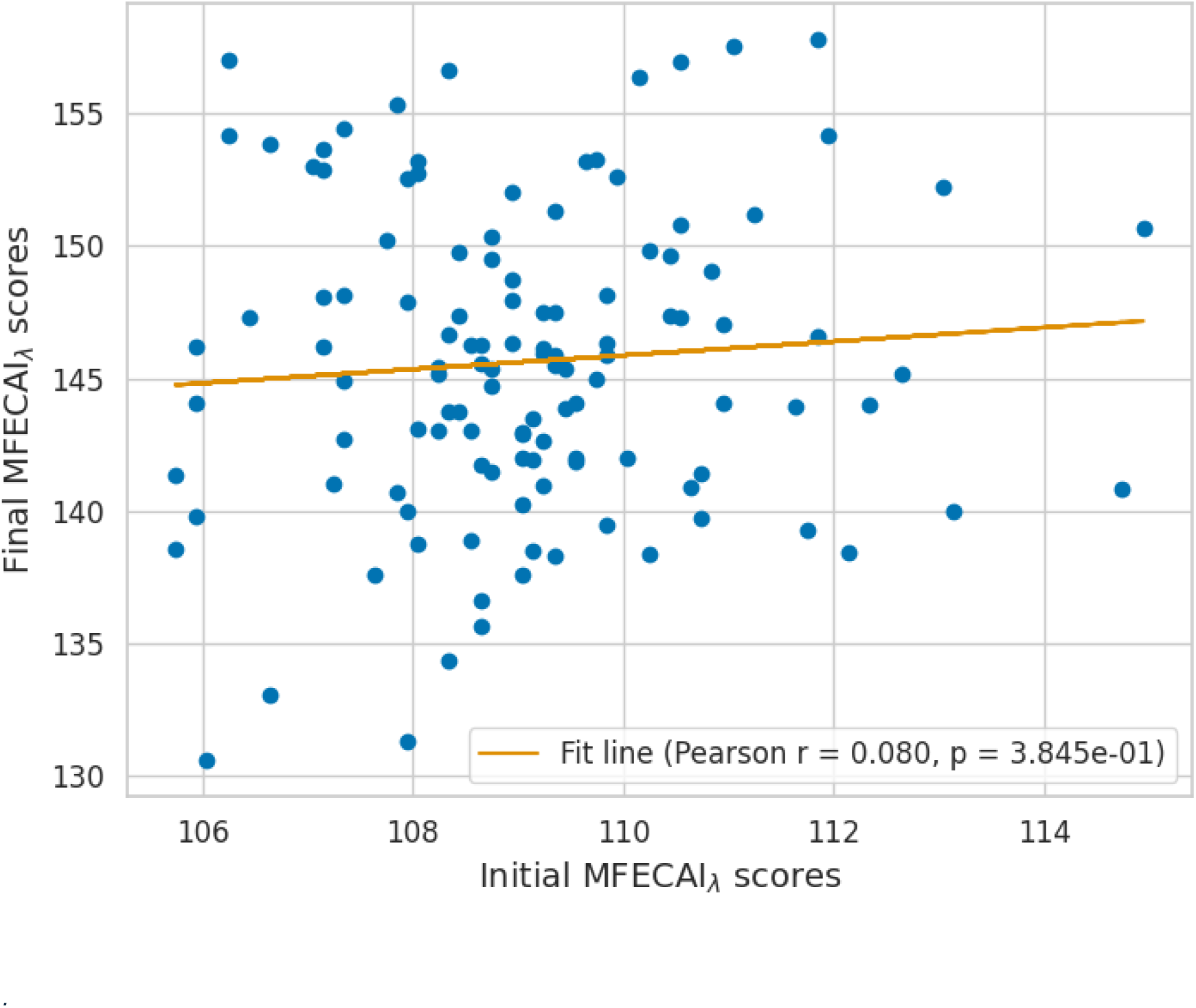
The correlation between Original and Final MFECAIλ values

**Figure S2.**
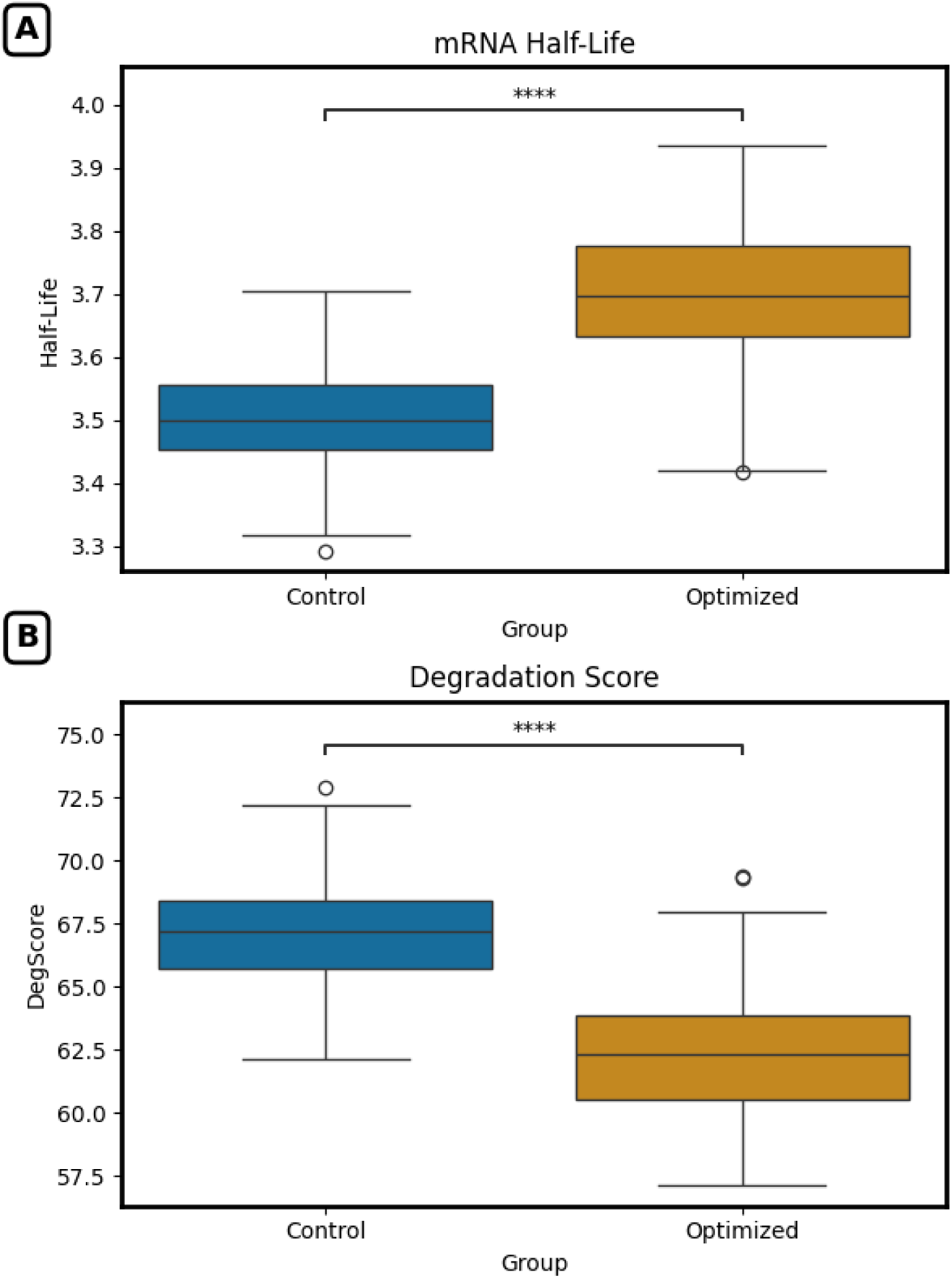
In-silico predictions of mRNA half-lives (in hours) and DeqScore values according to the DegScore model of Leppek et al. (2022). A: Predicted intracellular half-lives of mRNAs from control group and the best-of-three set of solutions. B: DegScore value for the same sequences as in (A). Higher scores indicate stronger propensity for degradation.

**Figure S3.**
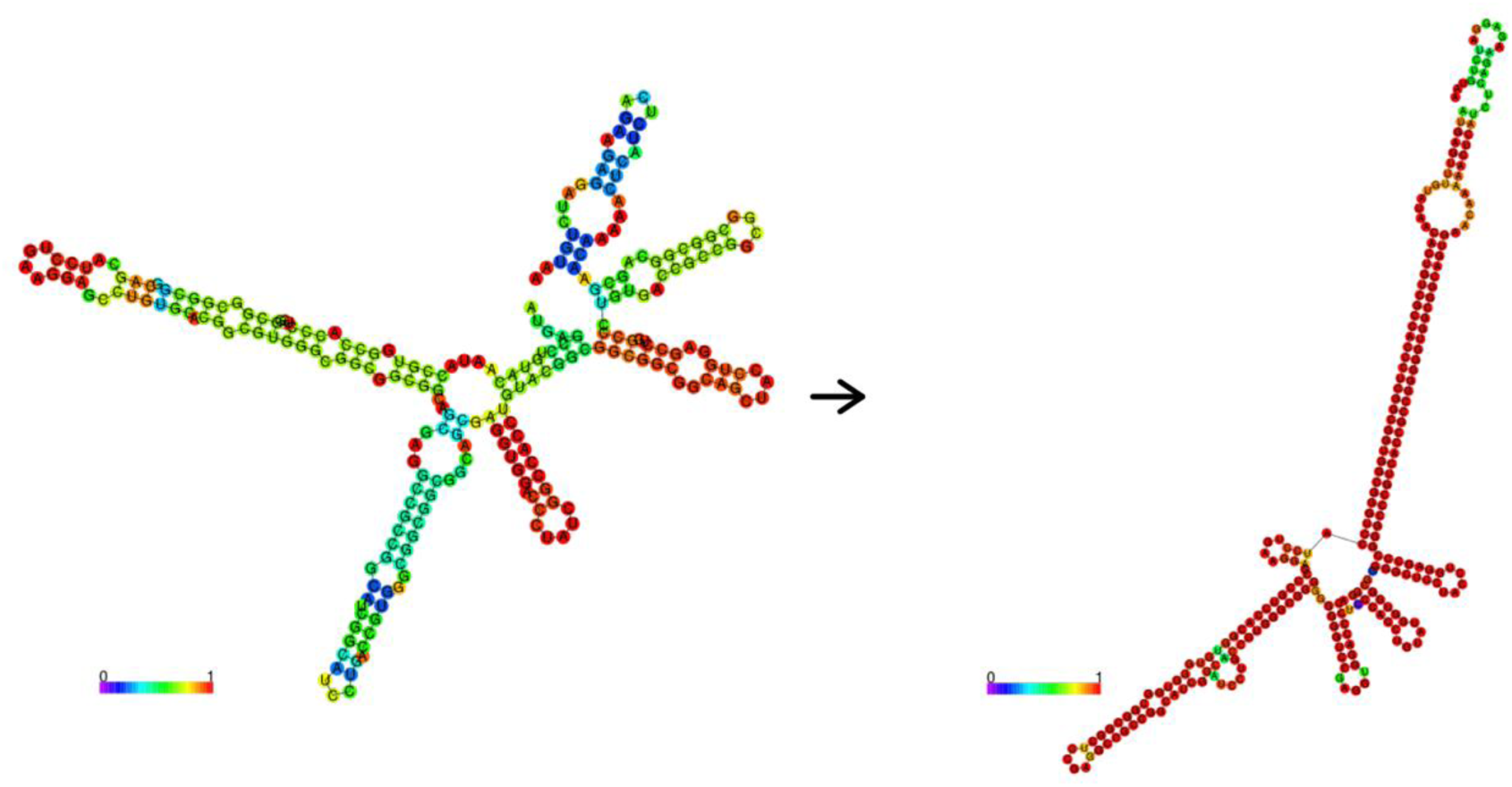
Secondary Structure comparison of the second best mRNA before and after sequence optimization.

**Figure S4.**
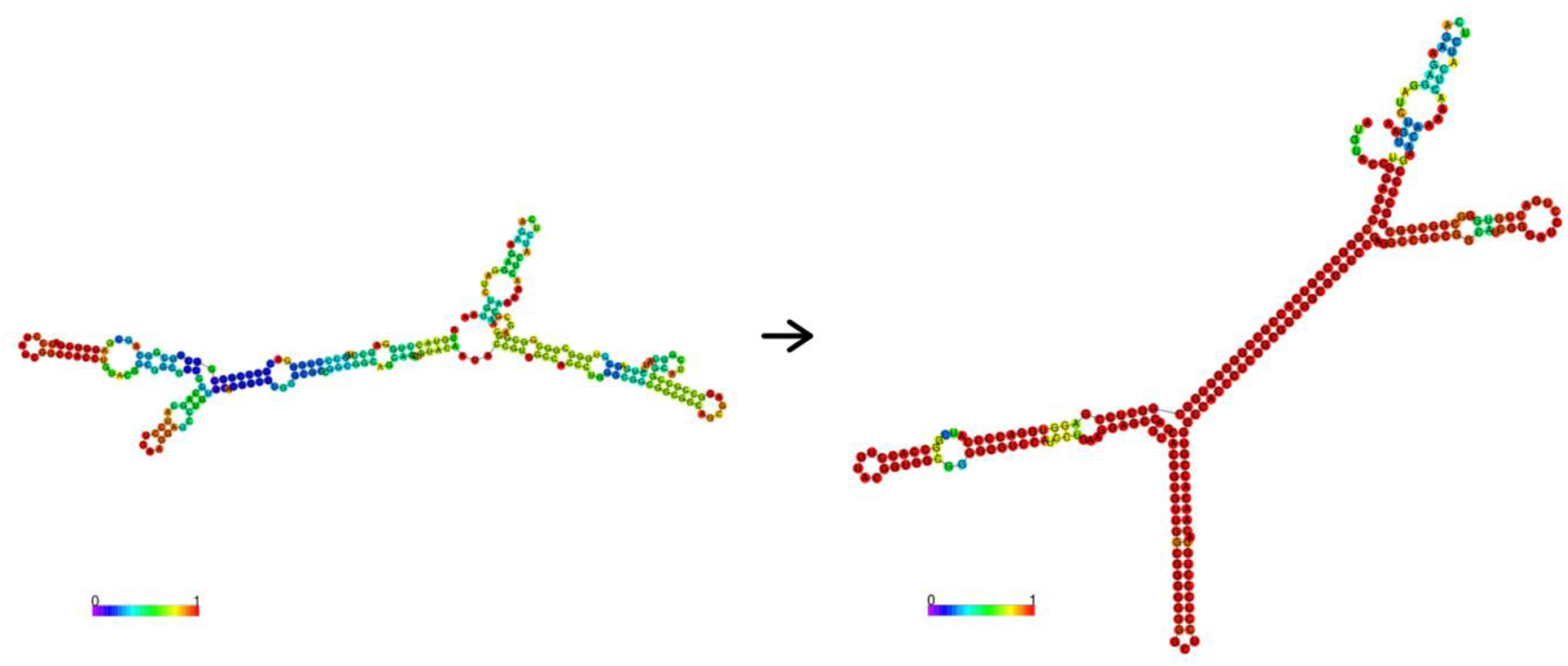
Secondary Structure comparison of the third best mRNA before and after sequence optimization.

**Table S1.** Grid search results: Top 5 hyperparameter value combinations according to MFECAIλ score(“Final Score”). The selected hyperparameters are highlighted in bold.

| **#** | **Learning rate** | **Batch Size** | **Hidden Size** | **Update rate** | **Average Final Score** | **Median Final Score** | **Time in seconds** |
|---|---|---|---|---|---|---|---|
| **1** | **0.01** | **64** | **128** | **10** | **141.93** | **142.03** | **232** |
| **2** | 0.01 | 64 | 256 | 10 | 141.69 | 141.54 | 669 |
| **3** | 0.01 | 96 | 256 | 10 | 141.42 | 141.54 | 714 |
| **4** | 0.01 | 128 | 256 | 10 | 140.63 | 140.55 | 730 |
| **5** | 0.01 | 96 | 128 | 10 | 140.15 | 140.09 | 296 |

**Table S2.** The peptides used for investigation of influence of sequence length on computation time.

| UniProt ID | Protein length |
|---|---|
| E9PCN6 | 59 |
| P13501 | 91 |
| Q99525 | 98 |
| P31949 | 105 |
| P0CE72 | 109 |
| P0CL80 | 117 |
| P01707 | 119 |
| O95867 | 125 |
| Q9GZQ8 | 125 |
| Q8NC38 | 126 |
| P15382 | 129 |
| Q9NV29 | 134 |
| O14880 | 152 |
| P54852 | 163 |
| Q14442 | 188 |
| Q15669 | 191 |

**Table S3.**
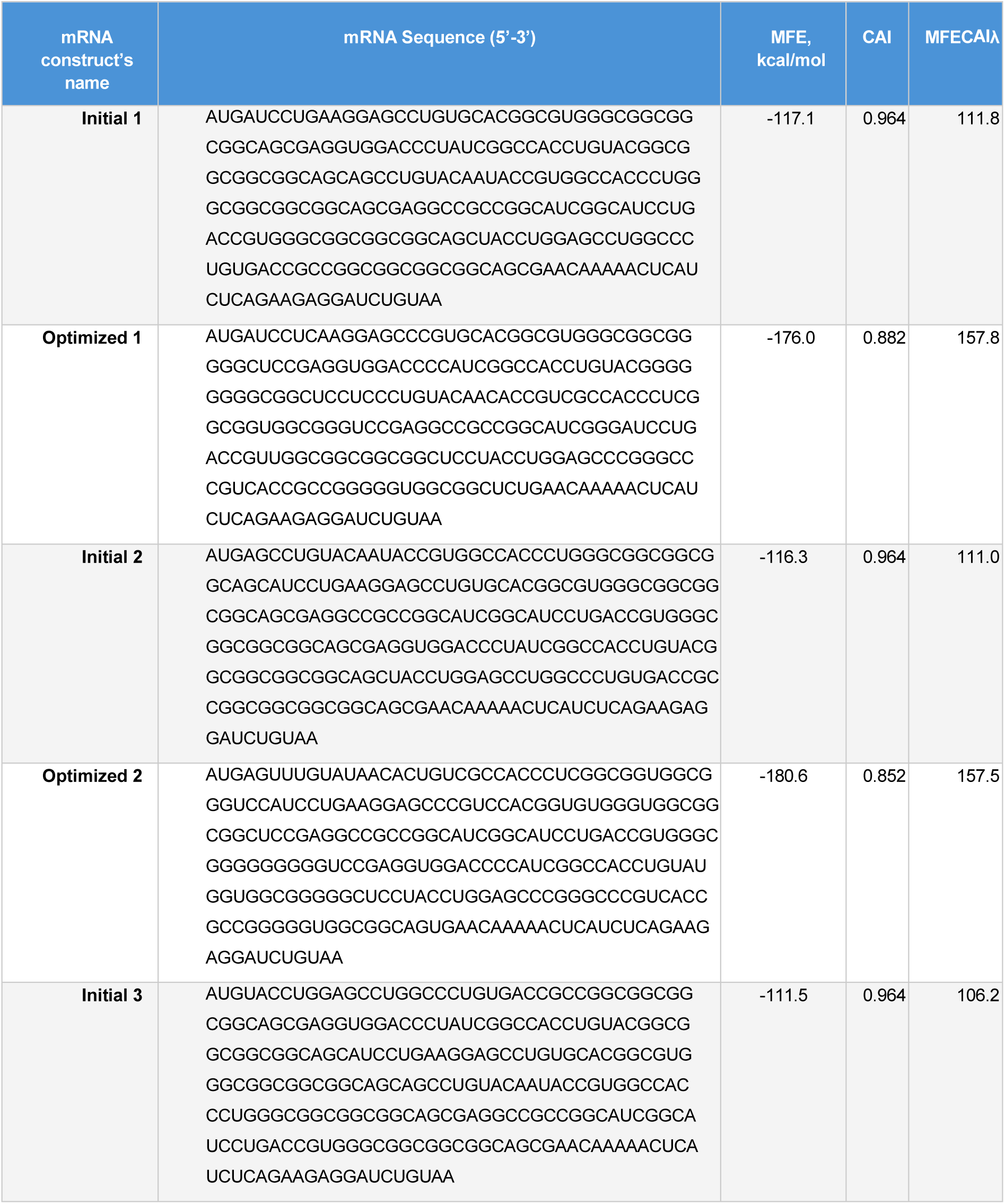
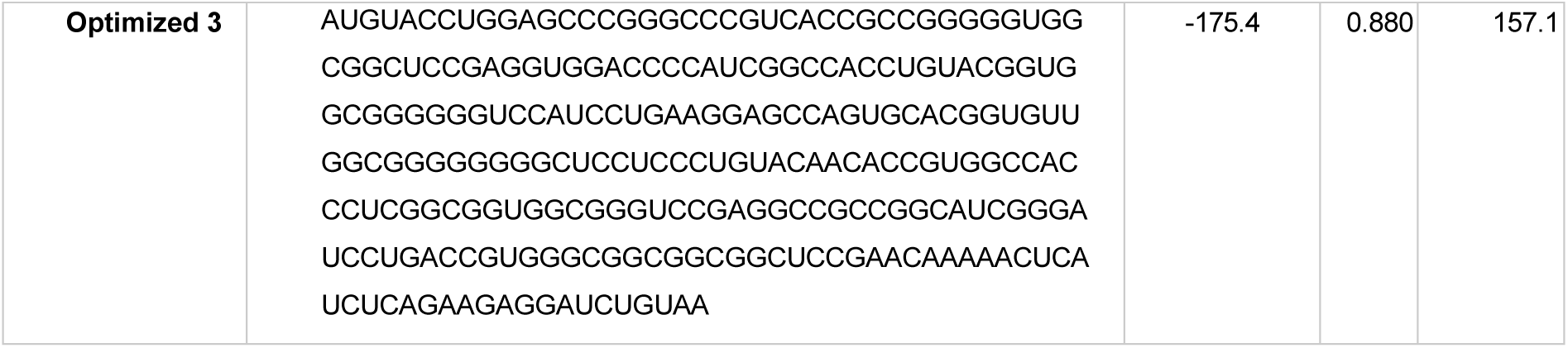
The mRNA sequences were selected for experimental validation. The optimal sequences are the top 3 solutions according to MFECAIλ given by ReinforSeqApp. Their corresponding initial sequences were selected as controls.

## Notes

https://github.com/JVeraLab/reinforseqpipe

### Competing Interest Statement

The authors have declared no competing interest.

## References

1. Amodu, O. A., Mahmood, R. A. R., Althumali, H., Bukar, U. A., Abdullah, N. F., & Jarray, C. (2025). Markov Decision Processes and Reinforcement Learning for Timely UAV-IoT Data Collection Applications (Vol. 1220). Springer Nature Switzerland. 10.1007/978-3-031-97011-5

2. Bahiri-Elitzur, S., & Tuller, T. (2021). Codon-based indices for modeling gene expression and transcript evolution. In Computational and Structural Biotechnology Journal (Vol. 19, pp. 2646–2663). Elsevier B.V. 10.1016/j.csbj.2021.04.042

3. Boyan, J. A., & Moore, A. W. (1995). Generalization in Reinforcement Learning: Safely Approximating the Value Function (Vol. 7). MIT Press.

4. Chaudhary, N., Weissman, D., & Whitehead, K. A. (2021). mRNA vaccines for infectious diseases: principles, delivery and clinical translation. In Nature Reviews Drug Discovery (Vol. 20, Number 11, pp. 817–838). Nature Research. 10.1038/s41573-021-00283-5

5. Cheng, F., Wang, Y., Bai, Y., Liang, Z., Mao, Q., Liu, D., Wu, X., & Xu, M. (2023). Research Advances on the Stability of mRNA Vaccines. In Viruses (Vol. 15, Number 3). MDPI. 10.3390/v15030668

6. Diament, A. (2022). *codon-bias (python package)*.

7. Fu H, Liang Y, Zhong X, Pan Z, Huang L, Zhang H, Xu Y, Zhou W, Liu Z (2020). Codon optimization with deep learning to enhance protein expression. In Sci Rep., 10(1):17617. 10.1038/s41598-020-74091-z.

8. Hanson, G., & Coller, J. (2018). Translation and Protein Quality Control: Codon optimality, bias and usage in translation and mRNA decay. In Nature Reviews Molecular Cell Biology (Vol. 19, Number 1, pp. 20–30). Nature Publishing Group. 10.1038/nrm.2017.91

9. Imdad, M. J., Khan, M. N., Alam, H. S., Khan, A. B., Mirani, Z. A., Khan, A., & Ahmed, F. (2023). Design and *in silico* analysis of mRNA vaccine construct against *Salmonella*. Journal of Biomolecular Structure and Dynamics, 41(15), 7248–7264. 10.1080/07391102.2022.2119280

10. Jain R, Jain A, Mauro E, LeShane K, Densmore D (2023). ICOR: improving codon optimization with recurrent neural networks. In BMC Bioinformatics, 24(1):132. 10.1186/s12859-023-05246-8.

11. Jin, L., Zhou, Y., Zhang, S., & Chen, S. J. (2025). mRNA vaccine sequence and structure design and optimization: Advances and challenges. In Journal of Biological Chemistry (Vol. 301, Number 1). American Society for Biochemistry and Molecular Biology Inc. 10.1016/j.jbc.2024.108015

12. Juszkiewicz, S., Slodkowicz, G., Lin, Z., Freire-Pritchett, P., Peak-Chew, S. Y., & Hegde, R. S. (2020). Ribosome collisions trigger cis-acting feedback inhibition of translation initiation. eLife, 9, 1–29. 10.7554/eLife.60038

13. Kames, J., Alexaki, A., Holcomb, D. D., Santana-Quintero, L. V., Athey, J. C., Hamasaki-Katagiri, N., Katneni, U., Golikov, A., Ibla, J. C., Bar, H., & Kimchi-Sarfaty, C. (2020). TissueCoCoPUTs: Novel Human Tissue-Specific Codon and Codon-Pair Usage Tables Based on Differential Tissue Gene Expression. Journal of Molecular Biology, 432(11), 3369–3378. 10.1016/j.jmb.2020.01.011

14. Kim, Y. A., Mousavi, K., Yazdi, A., Zwierzyna, M., Cardinali, M., Fox, D., Peel, T., Coller, J., Aggarwal, K., & Maruggi, G. (2024). Computational design of mRNA vaccines. Vaccine, 42(7), 1831–1840. 10.1016/j.vaccine.2023.07.024

15. Kozak, M. (1991). Structural features in eukaryotic mRNAs that modulate the initiation of translation. Journal of Biological Chemistry, 266(30), 19867–19870. 10.1016/s0021-9258(18)54860-2

16. Krueger, R., & Ward, M. (2024). Scalable Differentiable Folding for mRNA Design. 10.1101/2024.05.29.594436

17. Lee DSM, Park J, Kromer A, Baras A, Rader DJ, Ritchie MD, Ghanem LR, Barash Y (2021). Disrupting upstream translation in mRNAs is associated with human disease. In Nat Commun, 12(1):1515. 10.1038/s41467-021-21812-1.

18. Leppek, K., Byeon, G. W., Kladwang, W., Wayment-Steele, H. K., Kerr, C. H., Xu, A. F., Kim, D. S., Topkar, V. V., Choe, C., Rothschild, D., Tiu, G. C., Wellington-Oguri, R., Fujii, K., Sharma, E., Watkins, A. M., Nicol, J. J., Romano, J., Tunguz, B., Diaz, F., … Das, R. (2022). Combinatorial optimization of mRNA structure, stability, and translation for RNA-based therapeutics. Nature Communications, 13(1). 10.1038/s41467-022-28776-w

19. Lischer, C., Eberhardt, M., Flamann, C., Berges, J., Güse, E., Wessely, A., Weich, A., Retzlaff, J., Dörrie, J., Schaft, N., Wiesinger, M., März, J., Schuler-Thurner, B., Knorr, H., Gupta, S., Singh, K. P., Schuler, G., Heppt, M. V., Koch, E. A. T., … Vera, J. (2024). Gene network-based and ensemble modeling- based selection of tumor-associated antigens with a predicted low risk of tissue damage for targeted immunotherapy. Journal for ImmunoTherapy of Cancer, 12(5). 10.1136/jitc-2023-008104

20. Lorenz, R., Bernhart, S. H., Höner Zu Siederdissen, C., Tafer, H., Flamm, C., Stadler, P. F., & Hofacker, I. L. (2011). ViennaRNA Package *2.0*. http://www.tbi.univie.ac.at/RNA.

21. Mahmud, M., Kaiser, M. S., Hussain, A., & Vassanelli, S. (2018). Applications of Deep Learning and Reinforcement Learning to Biological Data. IEEE Transactions on Neural Networks and Learning Systems, 29(6), 2063–2079. 10.1109/TNNLS.2018.2790388

22. Malonis, R., Lai, J., & Vergnolle, O. (2020). Peptide-based vaccines. Chem Rev., 6(120), 3210–3229.

23. Mauger, D. M., Joseph Cabral, B., Presnyak, V., Su, S. V., Reid, D. W., Goodman, B., Link, K., Khatwani, N., Reynders, J., Moore, M. J., & McFadyen, I. J. (2019). mRNA structure regulates protein expression through changes in functional half-life. Proceedings of the National Academy of Sciences of the United States of America, 116(48), 24075–24083. 10.1073/pnas.1908052116

24. Mnih, V., Kavukcuoglu, K., Silver, D., Rusu, A. A., Veness, J., Bellemare, M. G., Graves, A., Riedmiller, M., Fidjeland, A. K., Ostrovski, G., Petersen, S., Beattie, C., Sadik, A., Antonoglou, I., King, H., Kumaran, D., Wierstra, D., Legg, S., & Hassabis, D. (2015). Human-level control through deep reinforcement learning. Nature, 518(7540), 529–533. 10.1038/nature14236

25. Nagarajan, P., Warnell, G., & Stone, P. (2019). Deterministic Implementations for Reproducibility in Deep Reinforcement Learning. http://arxiv.org/abs/1809.05676

26. Parhiz, H., Atochina-Vasserman, E. N., & Weissman, D. (2024). mRNA-based therapeutics: looking beyond COVID-19 vaccines. In The Lancet (Vol. 403, Number 10432, pp. 1192–1204). Elsevier B.V. 10.1016/S0140-6736(23)02444-3

27. Pfeifer, B. A., Beitelshees, M., Hill, A., Bassett, J., & Jones, C. H. (2023). Harnessing synthetic biology for advancing RNA therapeutics and vaccine design. In npj Systems Biology and Applications (Vol. 9, Number 1). Nature Research. 10.1038/s41540-023-00323-3

28. Romero, P., Banchereau, J., Bhardwaj, N., Cockett, M., Disis, M. L., Dranoff, G., Gilboa, E., Hammond, S. A., Hershberg, R., Korman, A. J., Kvistborg, P., Melief, C., Mellman, I., Palucka, A. K., Redchenko, I., Robins, H., Sallusto, F., Schenkelberg, T., Schoenberger, S., & Sosman, J. (2016). The Human Vaccines Project: A roadmap for cancer vaccine development. www.ScienceTranslationalMedicine.org

29. Roth, A. C. (2012). Decoding properties of tRNA leave a detectable signal in codon usage bias. Bioinformatics, 28(18). 10.1093/bioinformatics/bts403

30. Sauerer T, Albrecht L, Sievers NM, Gerer KF, Hoyer S, Dörrie J, Schaft N (2024). Electroporation of mRNA as a Universal Technology Platform to Transfect a Variety of Primary Cells with Antigens and Functional Proteins. In Methods Mol Biol., 2786:219–235. 10.1007/978-1-0716-3770-8_10.

31. Shahraki, P. K., Kiani, R., Siavash, M., & Bemani, P. (2025). Design of a multi-epitope vaccine against Staphylococcus Aureus lukotoxin ED using in silico approaches. Scientific Reports, 15(1). 10.1038/s41598-025-85147-3

32. Sharp, P. M., & Li, W.-H. (1987). The codon adaptation index-a measure of directional synonymous codon usage bias, and its potential applications. In Nucleic Acids Research (Vol. 15). https://academic.oup.com/nar/article/15/3/1281/1166844

33. Shen, W., Wang, D., Ye, B., Shi, M., Ma, L., Zhang, Y., & Zhao, Z. (2015). GC3-biased gene domains in mammalian genomes. Bioinformatics, 31(19), 3081–3084. 10.1093/bioinformatics/btv329

34. Terai, G., Kamegai, S., & Asai, K. (2016). CDSfold: An algorithm for designing a protein-coding sequence with the most stable secondary structure. Bioinformatics, 32(6), 828–834. 10.1093/bioinformatics/btv678

35. Tsitsiklis, J. N., & Van Roy, B. (1997). An Analysis of Temporal-Difference Learning with Function Approximation. In *IEEE* TRANSACTIONS ON AUTOMATIC CONTROL (Vol. 42, Number 5).

36. Van Houdt, G., Mosquera, C., & Nápoles, G. (2020). A review on the long short-term memory model. Artificial Intelligence Review, 53(8), 5929–5955. 10.1007/s10462-020-09838-1

37. Wayment-Steele, H. K., Kladwang, W., Watkins, A. M., Kim, D. S., Tunguz, B., Reade, W., Demkin, M., Romano, J., Wellington-Oguri, R., Nicol, J. J., Gao, J., Onodera, K., Fujikawa, K., Mao, H., Vandewiele, G., Tinti, M., Steenwinckel, B., Ito, T., Noumi, T., … Das, R. (2022). Deep learning models for predicting RNA degradation via dual crowdsourcing. Nature Machine Intelligence, 4(12), 1174–1184. 10.1038/s42256-022-00571-8

38. Wright, F. (1990). The “effective number of codons” used in a gene. Gene, 87(1), 23–29. 10.1016/0378-1119(90)90491-9

39. Wu, Z., Lou, J., Cheng, Y., Li, M., Wang, Z., Liu, N., Gao, X., Zheng, A., & Zhang, H. (2026). mRNA lipid nanoparticle vaccines: current status, challenges and future prospects. Molecular Biomedicine, 7(1), 186. 10.1186/s43556-026-00600-7

40. Xia, X. (2021). Detailed dissection and critical evaluation of the pfizer/biontech and moderna mrna vaccines. Vaccines, 9(7). 10.3390/vaccines9070734

41. Zhang, H., Liu, H., Xu, Y., Huang, H., Liu, Y., Wang, J., Qin, Y., Wang, H., Ma, L., Xun, Z., Hou, X., Lu, T. K., & Cao, J. (2025). Deep generative models design mRNA sequences with enhanced translational capacity and stability. Science. 10.1126/science.adr8470

42. Zhang, Zhang, L., Lin, A., Xu, C., Li, Z., Liu, K., Liu, B., Ma, X., Zhao, F., Jiang, H., Chen, C., Shen, H., Li, H., Mathews, D. H., Zhang, Y., & Huang, L. (2023). Algorithm for optimized mRNA design improves stability and immunogenicity. Nature, 621(7978), 396–403. 10.1038/s41586-023-06127-z

43. Zhao, P., Zhang, W.-B., & Chen, S.-J. (2010). Predicting Secondary Structural Folding Kinetics for Nucleic Acids. Biophysical Journal, 98(8), 1617–1625. 10.1016/j.bpj.2009.12.4319

44. Zuber, J., & Mathews, D. H. (2019). Estimating uncertainty in predicted folding free energy changes of RNA secondary structures. RNA, 25(6), 747–754. 10.1261/rna.069203.118

